# Cell networks in the mouse liver during partial hepatectomy

**DOI:** 10.1101/2023.07.16.549116

**Authors:** Bin Li, Daniel Rodrigo-Torres, Carl Pelz, Brendan Innes, Pamela Canaday, Sunghee Chai, Peter Zandstra, Gary D. Bader, Markus Grompe

## Abstract

In solid tissues homeostasis and regeneration after injury involve a complex interplay between many different cell types. The mammalian liver harbors numerous epithelial and non-epithelial cells and little is known about the global signaling networks that govern their interactions. To better understand the hepatic cell network, we isolated and purified 10 different cell populations from normal and regenerative mouse livers. Their transcriptomes were analyzed by bulk RNA-seq and a computational platform was used to analyze the cell-cell and ligand-receptor interactions among the 10 populations. Over 50,000 potential cell-cell interactions were found in both the ground state and after partial hepatectomy. Importantly, about half of these differed between the two states, indicating massive changes in the cell network during regeneration. Our study provides the first comprehensive database of potential cell-cell interactions in mammalian liver cell homeostasis and regeneration. With the help of this prediction model, we identified and validated two previously unknown signaling interactions involved in accelerating and delaying liver regeneration. Overall, we provide a novel platform for investigating autocrine/paracrine pathways in tissue regeneration, which can be adapted to other complex multicellular systems.

**Highlights:** A platform predicting cell-cell interactions in liver regeneration was established

This platform identified the BMP4 pathway antagonist Fstl1 as a stimulator of hepatocyte proliferation

This platform also discovered the role of Wnt pathway inhibitor Sfrp1 delaying liver regeneration

## Introduction

Rodent liver can fully restore itself to its former mass even after 70% partial hepatectomy (PHx) and this system represents a well-studied paradigm in regenerative medicine (Azuma et al., 2007). Hepatocytes, the liver parenchymal cells, quickly proliferate after PHx to replace the liver mass. After PHx, hepatocyte DNA synthesis peaks at 24 hours in rats and 36 hours in mice (Michalopoulos and DeFrances, 1997). Previous studies have examined the different cell types participating in PHx-induced liver regeneration. For example, liver sinusoidal endothelial cells (LSECs), one of the main cells in the hepatic cell niche, regulate liver regeneration and hepatocyte proliferation (Ding et al., 2010). In addition to LSECs, hepatic stellate cells (HSCs) are also involved in controlling hepatocyte proliferation (Michalopoulos, 2007).

Simultaneously, dividing hepatocytes produce paracrine signaling to activate other nonparenchymal cells (NPCs) (Michalopoulos and DeFrances, 1997) including LSECs, biliary epithelial cells (BECs), and Kupffer cells (KCs). KCs contribute to hepatocyte regeneration via TNF-α signaling (Shinozuka et al., 1994). While the contribution of individual liver cell types to liver regeneration has been previously examined by looking at their pairwise interactions with hepatocytes (Ding et al., 2010; Huch et al., 2013; Kordes et al., 2014; Li et al., 2017), a global view of all cell-cell interactions in the adult liver ground state and their changes during regeneration has not been available to date. Importantly, many other cell types, including cholangiocytes and endothelial cells, also have to divide after partial hepatectomy to fully restore the tissue. Very little is known about the signaling events that govern the regeneration of non-hepatocytes.

Cells communicate with each other by ligand-receptor interactions via paracrine and autocrine pathways to initiate the responses to partial hepatectomy. One well-studied example is hepatocyte growth factor (HGF), a ligand secreted by HSC and LSEC, which interacts with the receptor c-Met on hepatocytes to induce hepatocyte proliferation. In rats, plasma HGF increases dramatically one hour after PHx (Michalopoulos and DeFrances, 1997). HGF overexpression *in vivo* induces homeostatic hepatocytes to enter G_0_/S phase and mitosis (Michalopoulos, 2007). Although several specific signaling pathways have been analyzed in detail based on a candidate molecule approach, an unbiased method for the identification of all possible signaling events governing liver regeneration has not been available. Given the thousands of potential ligands and receptors in the liver, it is likely that many functionally important signaling pathways remain to be discovered.

Previous reports from the Zandstra lab described the structure and dynamics of the hierarchically organized blood system at the cell-cell interaction and intramolecular network levels (Kirouac et al., 2009; Kirouac et al., 2010). More r ecently, a significantly improved bioinformatics platform was developed for constructing connectivity-based intercellular signaling networks using the upregulated ligand and receptor genes of different cell types in a tissue (Kirouac et al., 2010). In particular, they explored the intercellular signaling from 11 human bone marrow hematopoietic populations (stem cells, erythroid progenitors, myeloid progenitors etc.). By integrating high-throughput molecular profiling (transcriptome and proteome), protein interaction and information, and mechanistic modeling with cell culture experiments, they showed that complex intercellular communication networks by secreted factors mediated intercellular communication networks and regulate blood stem cell fate decisions.

Several novel unknown ligand/receptor interactions were discovered and validated in HSC expansion cultures *in vitro*. Here, we adapted this platform to a liver regeneration system to define autocrine and paracrine cell interactions and to predict novel regulators of tissue behavior. We isolated 10 liver cell populations from homeostatic adult mice and after induction of liver regeneration by partial hepatectomy. We performed bulk RNA sequencing (RNA-seq) independently on each population and constructed cell-cell interactions (CCInx) (Ximerakis et al., 2019) by matching the ligand and receptor pairs in the ligand-receptor database. A dense map of potential intercellular interactions was identified and two previously unknown receptor-ligand interactions important for liver regeneration were discovered and functionally validated.

## Results

### Isolation of liver cells with surface markers

To isolate liver cells, we performed a two-step collagenase perfusion of C57B/L6 wildtype male mice liver at 8 weeks in the normal state and 24 hrs after 70% PHx. Hepatocytes and NPCs were collected and labelled with antibodies according to published papers (Table 1) (Alaverdi, 2004; Ding et al., 2010; Dorrell et al., 2011; Hoppo et al., 2004; Huch et al., 2013; Kumar et al., 2006; Li et al., 2017; Mederacke et al., 2015). FACS was used to isolate 10 distinct populations of interest: Regular biliary epithelial cells (ST14^-^CD26^-^MIC1-1C3^+^CD31^-^CD45^-^CD11b^-^, n = 4 in homeostasis (Li et al., 2017), n = 2 in PHx), blood cells (CD45^+^, Figure S1A, n = 2 in homeostasis, n = 2 in PHx) and Thy1+ (Thy1^+^CD45^-^, Figure S1A, n = 2 in homeostasis, n = 2 in PHx) cells., clonogenic biliary epithelial cells (ST14^+^CD26^-^MIC1-1C3^+^CD31^-^CD45^-^CD11b^-^, n = 4 in homeostasis(Li et al., 2017), n = 2 in PHx), endothelial cells (Lyve1^-^ CD34^+^CD144^+^CD309^+^CD45^-^, Figure S1B, n = 2 in homeostasis, n = 3 in PHx), liver sinusoidal endothelial cells (Lyve1^+^CD34^-^CD144^+^CD309^+^CD45^-^, Figure S1B, n = 3 in homeostasis, n = 5 in PHx), hepatocytes (HCs) (OC2-2F8^+^CD45^-^CD31^-^, Figure S1C, n = 4 in homeostasis, n = 4 in PHx), hepatic stellate cells (CD146^+^CD45^-^CD31^-^Violet^+^, Figure S1D, n = 2 in homeostasis, n = 3 in PHx), Kupffer cells (F4/80^+^CD11b^+^, Figure S1E, n = 2 in homeostasis, n = 3 in PHx), non-progenitor ducts (NPD) (CD26^+^MIC11C3^+^CD31^-^CD45^-^CD11b^-^, n = 2 in homeostasis, n = 2 in PHx). At least 10,000 cells were collected for each population to extract bulk RNA.

**Table 1.**
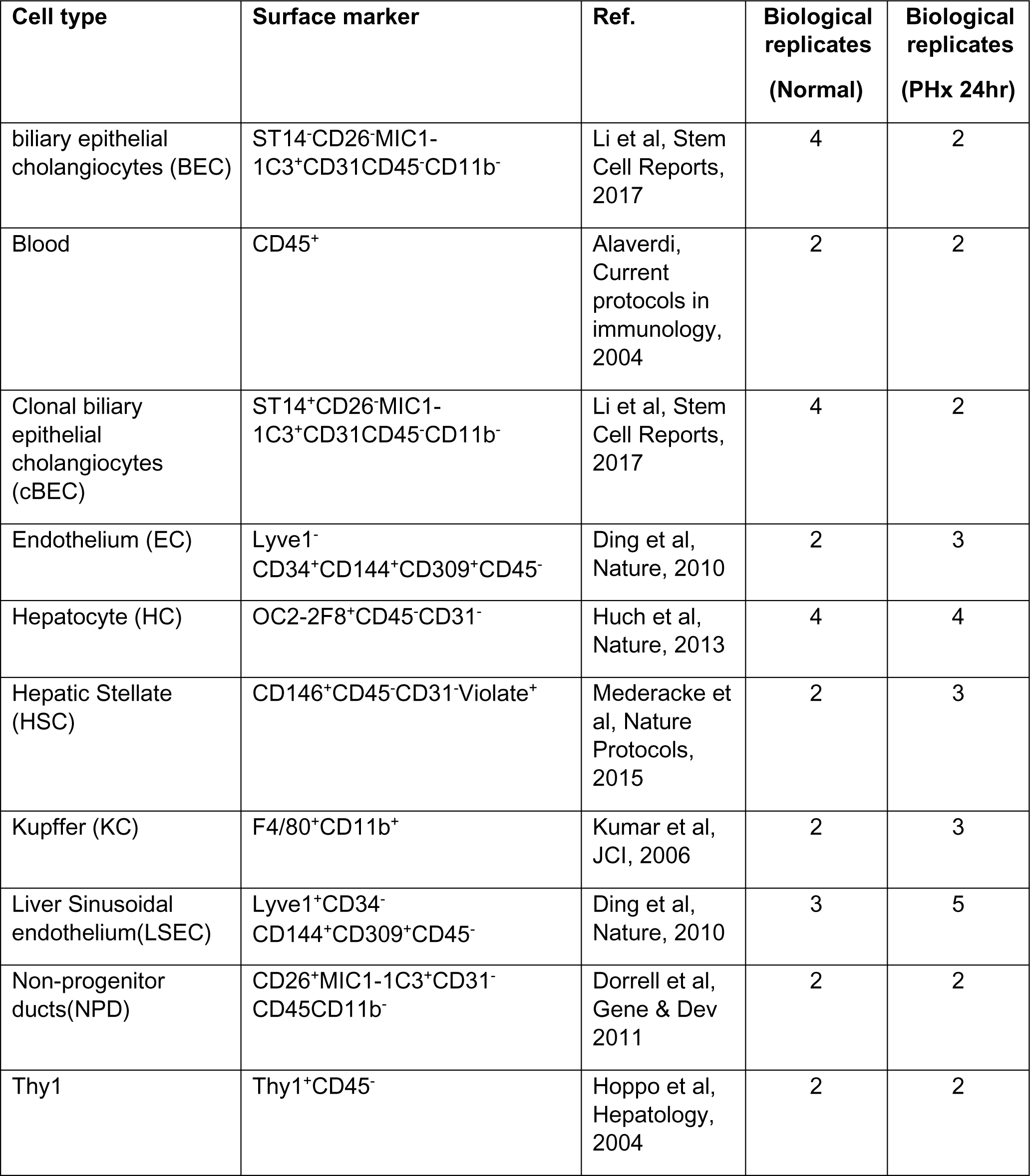
Cell isolation strategy.

### Transcriptome and cell-cell interaction analysis in mouse liver cells

To compare these populations at the transcriptional level, we extracted RNA from freshly FACS-sorted cells to perform RNA-seq. The data were analyzed using a previously published computer algorithm (Qiao et al., 2014) with a substantially expanded cell interaction database (Ximerakis et al., 2019). We found 39,630 common cell interactions (including autocrine and paracrine pathways) among the 10 cell types that were present in both normal and PHx liver (Figure 1A, Table S1 and S2). In addition, we identified 19,528 global interactions that were unique to the ground state, whereas there were 23,640 global interactions present only after PHx (Figure 1A, Table S1 and S2). We next narrowed our analysis to look at only those interactions that involved hepatocytes. Hepatocytes participated in 6,645 interactions, 8.0% of the total (Figure 1B, Table S1 and S2). 3,342 were present in both homeostasis and the regenerative state. In addition, there were 1,632 unique ground state interactions (8.4% of global interactions) and 1,671 interactions present only after PHx (7.1% of global interactions) (Figure 1B, Table S1 and S2). To get a sense of how the cell network changes during regeneration, we looked at the top 10 ranked ligand-receptor interactions (based on CCInx weights calculated using our log2 FC calculations for both ligand and receptor) for each cell type in both the normal state (Figure 1C, Table S3) and after PHx (Figure 1D, Table S4). It is immediately apparent that the network changes between the two states and that input and outputs are different for all cell types. Interestingly, the majority of predicted top interactions occurred between non-hepatocyte populations (Figure 1C, D, Table S3 and S4). Among the top 73 interactions in normal livers, there was only 1 paracrine pathway coming from another cell type (NPD: ligand C3) to hepatocytes (receptor: Cfi) but there were 22 outward paracrine pathways from hepatocytes to other cell types (2 to BEC, 2 to cBEC, 3 to NPD, 4 to EC, 4 to LSEC, 0 to blood, 3 to Thy1, 0 to KC, 4 to HSC) (Figure 1C, Table S3). In PHx liver, the number of signals from hepatocytes->others and others->hepatocytes were 7 (1 to LSEC,1 to Thy1, 1 to KC, 4 to HSC) and 3 (HSC: ligand Ecm1, receptor Cfi, NPD: ligand C3, receptor Cfi, Thy1: ligand C3, receptor Cfi), respectively (Figure 1D, Table S4).

**Figure 1.**
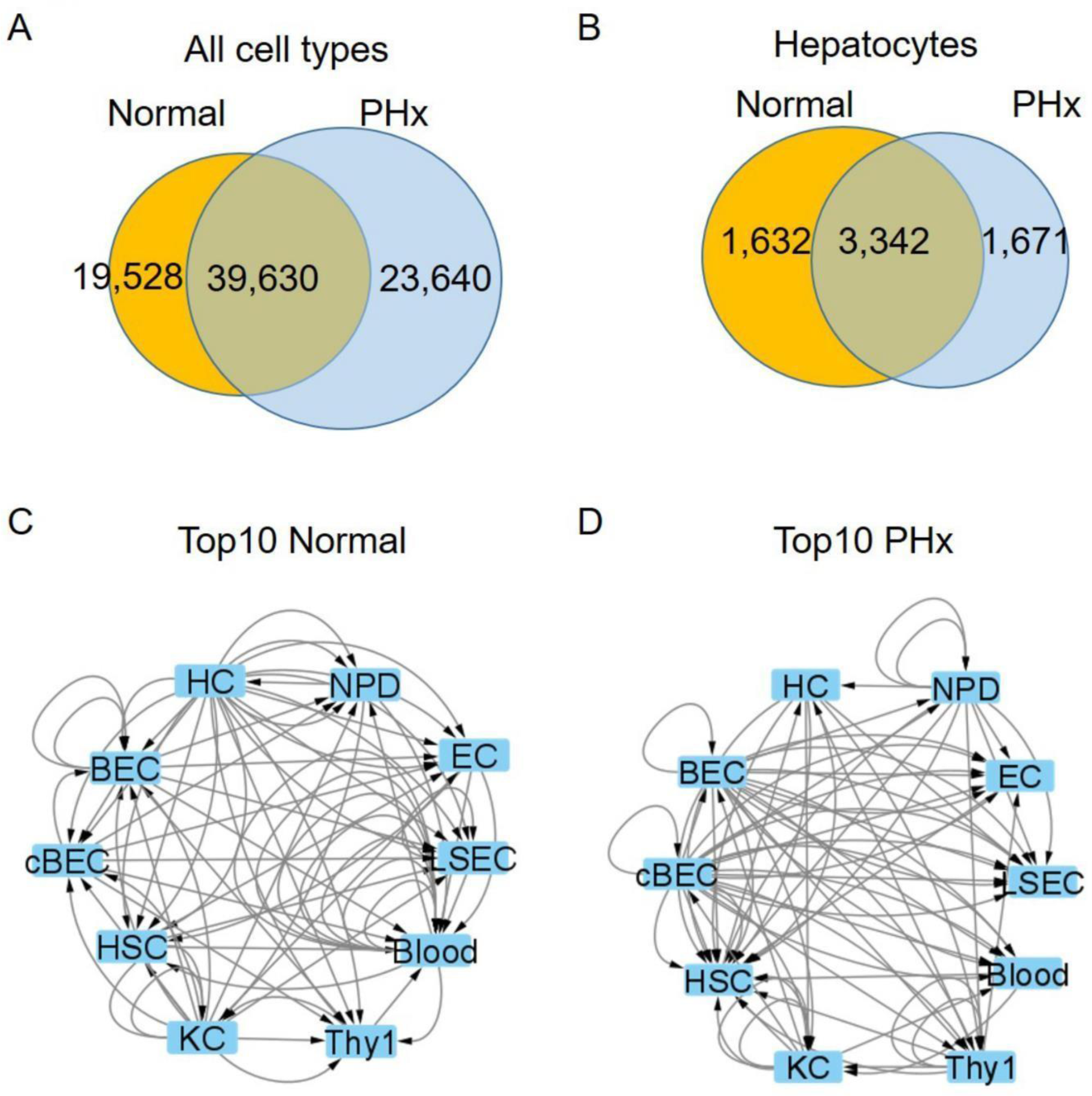
Cell-cell Interactions in homeostatic and regenerative liver. (A-B) Venn diagram of the number of paracrine and autocrine interactions in normal and partial hepatectomy (PHx) among all 10 cell types (A) and hepatocytes only (B). (A) There are 19,528 unique interactions in normal liver and 23,640 in PHx. The normal and PHx states had 39,630 shared common interactions. (B) In hepatocytes, there were 1,632 unique interactions in normal and 1,671 after PHx, with 3,342 shared common interactions. (C-D) Schematic top 10 CCInx weighed interactions in normal (C) and PHx (D) livers by Cytoscape network visualization. The interactions are ranked by the edge weights calculated by CCInx. PHx: 24 hr after 70% partial hepatectomy. HC: hepatocyte. NPD: non-progenitor duct cells. EC: Endothelial cells. LSEC: liver sinusoidal endothelial cells. Blood: blood lineage cells. Thy1: Thy1+ cells. KC: Kupffer cell. HSC: Hepatic Stellate cell. BEC: biliary epithelial cell. cBEC: clonal biliary epithelial cell. Arrows indicate either autocrine (from one cell to itself) or paracrine (from one cell to another) interactions.

### Validation of known ligands by CCInx

To determine whether our system could capture transcriptionally regulated receptor-ligand pairs that are known to be important in liver regeneration, we examined such molecules using the previously reported cell-cell interaction database (CCInx) (Ximerakis et al., 2019). It has been reported that Pdgfβ and Tgfα induce hepatocyte proliferation both *in vivo* and *in vitro*, respectively (Li et al., 2022; Mead and Fausto, 1989; Vrochides et al., 1996). Our cell network platform verified that Pdgfβ paracrine and autocrine pathways were “off” in homeostasis (Figure S2A) while “on” after PHx (Figure S2B). We also investigated the Tgfα pathway in homeostasis and PHx in our database (Figure S2C and D). As expected, the Tgfα pathway was inactive in LSECs during homeostasis (Figure S2C) but activated after PHx (Figure S2D).

### Fstl1 as a novel ligand in accelerating liver regeneration

In addition to known pathways, our analysis revealed many previously unexplored receptor-ligand interactions that changed during regeneration and were therefore candidates to play a functional role. To probe the predictive value of our network, we performed functional tests on two of these candidates. We mined the data to identify novel candidate ligands that are only present and highly expressed in either homeostasis or perturbation (regenerative liver). A literature review helped us to determine that these molecules had not been previously assessed in the context of liver regeneration. Based on these criteria, we chose the ligands follistatin like 1 (Fstl1) and secreted frizzled related protein 1 (Sfrp1) for functional assays. Fstl1 interactions from sinusoidal endothelial cells to hepatocytes were “off” in homeostasis (Figure 2A, 2C and 2E) but on after 24 hr PHx (Figure 2B, 2D and 2E). In addition, Fstl1 expression in homeostatic LSECs was inactivated but was strongly upregulated in PHx (Figure 2E).

**Figure 2.**
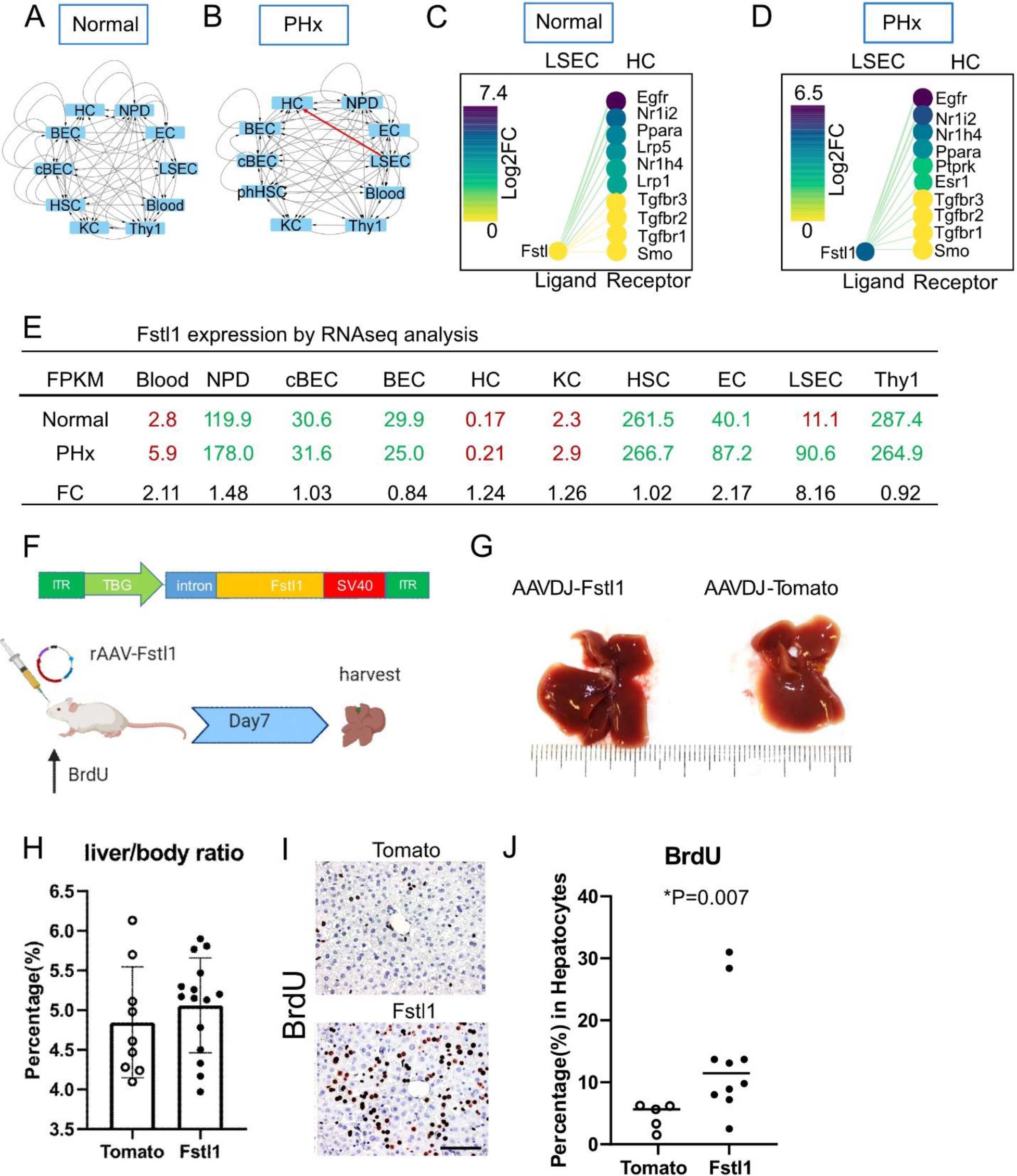
Fstl1 overexpression promotes hepatocyte proliferation. (A-B) Cytoscape network visualization of Fstl1 as a ligand in 10 different cell types from the normal (A) and PHx (B) liver. Arrows indicate the paracrine and autocrine signal pathways. Red arrow indicates the paracrine signal pathway appears in the PHx but not in the normal liver. (C-D) The CCInx database showed Fstl1 as a ligand in LSEC and its top 10 receptors in HC. Fstl1 is off (yellow circle) (C) in the normal liver, but upregulated and on (dark purple circle) after partial hepatectomy (D). log2 FC: log2 fold change. (E) Normalized gene expression of *Fstl1* (Tag counts in FPKM) in 10 different cell types from normal liver and PHx. Tag counts in red color: the *Fstl1* gene is “off”; Tag counts in green color: the *Fstl1* gene is “on”. FC: fold change. (F) rAAV construct for overexpressing the *Fstl1* gene. The *Fstl1* gene was cloned into a AAV2 vector backbone with the human thyroid hormone-binding globulin (TBG) as promoter. 2×10^11^ vector genomes of AAV-Fstl1 virus were injected into C57BL/6 mice fed with BrdU in the drinking water. The liver was harvested on day 7 after injection. (G) Representative morphology of mouse liver after Fstl1 overexpression. A liver transduced with an AAVDJ-Tomato vector was used as control. (H) Liver/body weight ratio of independent mice treated with AAVDJ-Tomato (n = 9) and AAVDJ-Fstl1 (n = 15). Student’s t test. P > 0.05 (I) Histology of anti-BrdU staining in the control (Tomato) and Fstl1 overexpression liver. Scale bar = 50 µm. BrdU expressing nuclei stained brown. (J) Percentage of BrdU+ hepatocytes from Fstl1 overexpression (n = 10) and mock (Tomato) (n= 5) transduced mouse liver. Student’s t test. *P = 0.007 LSEC: liver sinusoidal endothelial cells. HC: hepatocytes

We then investigated the expression levels of Fstl1 in all different cell types (Figure 2E). Blood cells, HCs and KCs had low FPKM (<10) before (FPKM = 2.8, 0.17 and 2.3 respectively) and after PHx (FPKM = 5.9, 0.21 and 2.9 respectively), suggesting the gene was completely “off” in these cells. Bile duct lineage cells BECs, cBECs, NPD, HSC and Thy1 populations had higher levels of Fstl1 but no significant change before and after PHx (Figure 2E). Endothelial cells (ECs and LSECs) were the only two populations that showed elevated Fstl1 expression after PHx. The physical proximity of LSECs to hepatocytes makes them of particular interest in terms of providing important ligands. Notably, compared to ECs, LSECs had a higher fold change (FC = 8.16) from homeostasis (FPKM = 11.1) to PHx (FPKM = 90.6), while EC had a lower increase (FC = 2.17) from homeostasis (FPKM = 40.1) to PHx (FPKM = 87.2).

To test the role of Fslt1 by gain of function *in vivo*, we constructed self-complementary recombinant adeno-associated virus (scrAAV) carrying the mouse *Fstl1* transgene (Figure 2F) and overexpressed Fstl1 in the liver. BrdU was added to the drinking water and its incorporation into nuclei was used to measure liver cell division over that time period. Seven days after treatment with rAAV-Fslt1, the livers were harvested (Figure 2F). Compared to control mice, in which rAAV carrying the tdTomato transgene was administrated, the liver size did not change (Figure 2G) macroscopically, nor did the liver/body weight ratio (AAV-Fstl1 5.1 ± 0.6% vs AAV-tdTomato 4.8 ± 0.7%, Figure 2H).

However, immunohistochemistry showed that after Fstl1 overexpression, >10% of hepatocytes were BrdU^+^ (14.1 ± 9.2%, n = 10 mice, Figure 2I and J). In contrast, the proportion of BrdU^+^ hepatocytes in AAV-tdTomato control was much lower (4.6 ± 2.1%, n = 5 mice, *p = 0.007, Figure 2J). No hepatic injury was seen by histology. However, when we add recombinant Fstl1 in the hepatocyte culture media, there was no significant hepatocyte proliferation (Figure S3). Together, these results show that Fstl1 can powerfully induce hepatocyte replication *in vivo* but not in vitro.

### Sfrp1 as a novel ligand for delaying liver regeneration

Using the same selection criteria, we identified another gene, secreted frizzled related protein 1 (Sfrp1), which is an antagonist of the Wnt signal pathway. Wnt/β-catenin signals drive PHx induced liver regeneration (Russell and Monga, 2018). We found that Sfrp1 interactions were active (14 interactions) during mouse liver homeostasis (Figure 3A and C) but were quiescent (0 interactions) in the PHx liver (Figure 3B and D). Among the 10 liver cell populations, Sfrp1 was exclusively expressed in Thy1 cells (FPKM > 10, Figure 3E). We observed that in the Thy1 population, Sfrp1 expression was significantly downregulated (FC = 0.19) from normal (FPKM = 60.4) to PHx (FPKM = 11.5).

**Figure 3.**
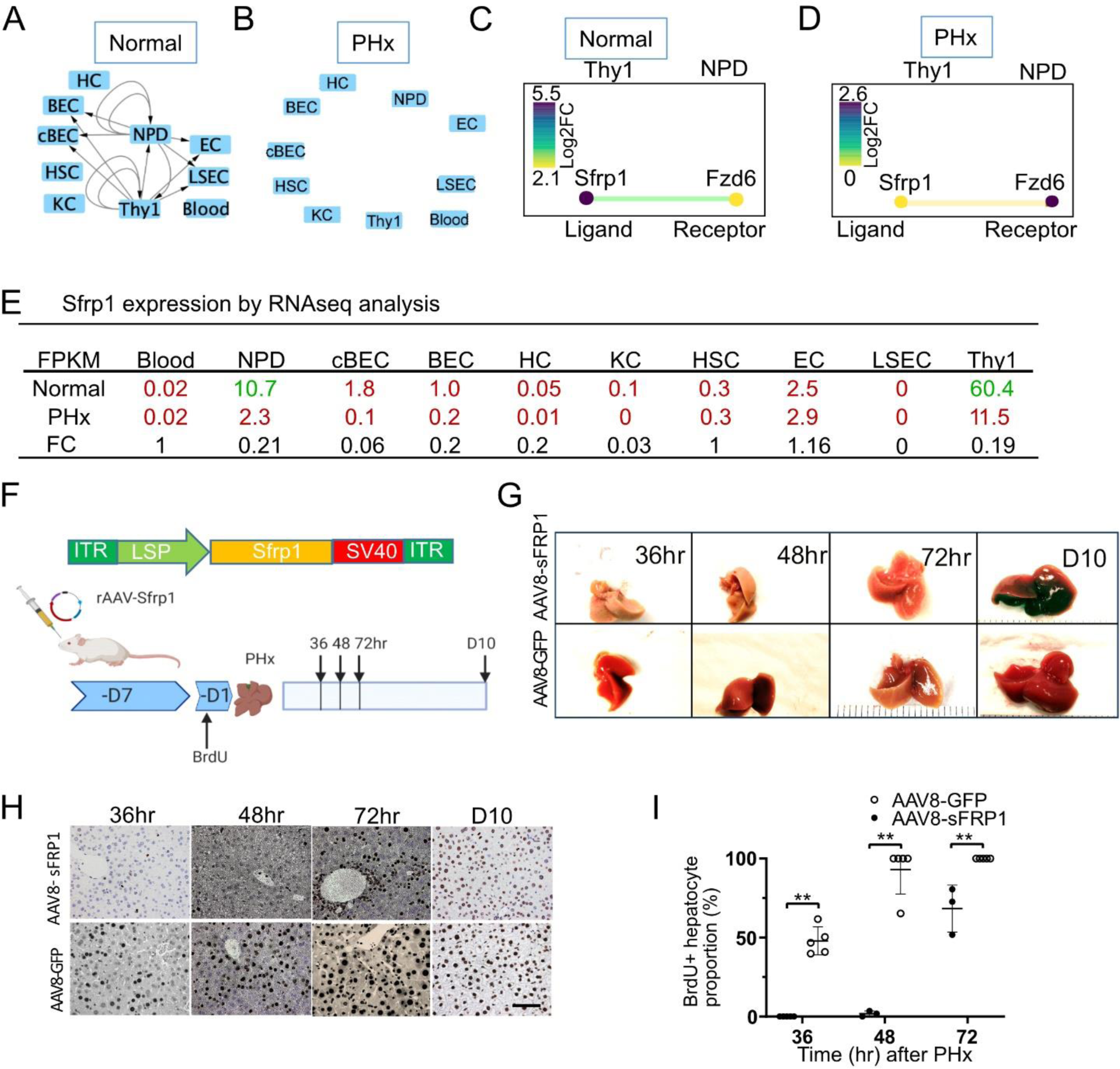
Sfrp1 delays liver regeneration. (A-B) Sfrp1 pathway in 10 different cells in the normal liver (A) and 24 hr after 70% partial hepatectomy (PHx) (B). Arrows indicate the paracrine and autocrine signal pathways. (C-D) The CCInx database showed Sfrp1 as a ligand in Thy1+ cells and its receptor Fzd6 in NPD. Log2 FC: log2 fold change. (E) Fragment per kilobase per million (FPKM) mapped reads in 10 different cell types from normal liver and PHx. Tag counts in red color: the *Sfrp1* gene is “off”; Tag counts in green color: the *Sfrp1* gene is “on”. FC: fold change. (F) AAV8-Sfrp1 was transfected into a wild type mouse 7 days before feeding with BrdU drinking water. One day after BrdU water treatment, partial hepatectomy was performed. Liver harvest was carried out on 36hr, 48hr, 72hr and 240hr (Day10) after partial hepatectomy. (G) Representative morphology of mouse liver overexpressing with Sfrp1 with partial hepatectomy at 36hr, 48hr, 72hr and 240hr (day10). Liver transfected with AAV8-GFP was used as control. (H) Histology of anti-BrdU staining in the mouse liver treated control (AAV8-GFP) and Sfrp1 over expressed mouse liver at different time points. Scale bar = 50 µm. BrdU expressing nuclei stained brown. (I) Quantitative assay of the percentage of Brdu^+^ hepatocytes in the mouse liver treated with control (AAV8-GFP, n = 3) and AAV8-Sfrp1(n = 5) at different time points. Statistical analyses: Mann-Whitney U test. **p< 0.01.

Thus, to evaluate the effects of gain-of Sfrp1 function *in vivo*, we constructed an rAAV carrying a Sfrp1 transgene (Figure 3F). rAAV-Sfrp1 was administrated 7 days before BrdU administration. One day after BrdU was added to the drinking water, PHx was performed to trigger liver regeneration (Figure 3F). We found that in the Sfrp1 overexpression cohort, the hepatocytes were not marked by BrdU until PHx 48 hr, whereas in the control group with rAAV-GFP only, hepatocytes incorporated BrdU as early as 36 hr after PHx (Figure 3H). Quantitative analysis demonstrated that after Sfrp1 overexpression, the percentage of BrdU^+^ hepatocytes was significantly lower than that in the AAV-GFP control at 36, 48 and 72 hr after PHx (Figure 3I). These results showed that Sfrp1 is a negative regulator of liver regeneration.

## Discussion

Our work described herein provides a first generation cell-cell interaction map based on transcriptome changes during liver regeneration. It is very clear from published work that liver regeneration represents a very complex reorganization of the structure of the entire organ and involves all the many cell types present. Past studies have been largely hepatocyte-centric and much valuable knowledge has been garnered from studying transcriptomic changes in whole liver RNA during regeneration. About 80% of total RNA in the liver comes from hepatocytes (Li et al., 2017; MacParland et al., 2018) and hence it is easy to detect RNA expression changes in this cell type. However, it is well known that non-hepatocyte liver cells are also very important in initiating and orchestrating liver regeneration (Ding et al., 2010; Huch et al., 2013; Kordes et al., 2014; Li et al., 2017). Genomic studies targeted at specific other cell populations have been published. An example is the well-known effect of angiocrine factors produced by sinusoidal endothelium (Ding et al., 2010). Based on the understanding that all liver cells have to regenerate to restore liver mass after partial hepatectomy we hypothesized that “all cells in the liver talk to all other cells” during the regenerative process. We therefore decided to apply the cell interaction network methodology that was successfully applied to the hematopoietic system to the liver. Our analysis provides a comprehensive database of transcriptomic changes in ten antigenically defined liver cell types. A very rich and dense network of potential interactions was revealed. Indeed, highly significant gene expression changes were observed in all ten cell types analyzed, confirming that regeneration requires adaptive changes in all liver resident cells.

It is important to emphasize that all of the interactions described here in our network represent only potential interactions. Transcriptome data alone do not prove that the ligands and receptors present at the RNA level in different cell types functionally interact. Our network serves to generate hypotheses that need to be validated empirically. Conversely, however, it is relatively safe to assume that interactions involving genes encoding ligands or receptors that are not expressed in a given cell type, are in fact not active in that cell. Together with information about the expression level of the ligands and receptors as well as literature information, these criteria can be used to prioritize the potential interactions for experimental testing, as we did here.

The analysis of mRNA alone represents a limitation of our method. There are well known signaling events during liver regeneration (Huh et al., 2004; Ishii et al., 1995; Kaibori et al., 2002; Nejak-Bowen et al., 2013; Patijn et al., 1998) that do not involve changes in the mRNA expression levels of ligands or their receptors. Signaling by HGF, for example, occurs very rapidly without transcriptional activation. Despite this obvious gap in our network, we believe that many important interactions are mirrored at the RNA level and that our network database will be useful to liver biology investigators.

To probe whether our network has any real life validity, we mined the data for yet unpublished cell-cell interactions and performed functional validation experiments with two of these. In both of the examples chosen, our gain-of-function experiments confirmed the activity of the candidate molecules in partial hepatectomy. We identified Fstl1 as a ligand produced by endothelial cells which significantly induces hepatocyte mitosis. FSTL1, a secreted glycoprotein that acts as a bone morphogenetic protein 4 (BMP4) antagonist (Geng et al., 2011), is reported to drive cardiomyocytes to enter the cell cycle in mice (Wei et al., 2015). Here, gain-of Fstl1 function *in vivo* also induced hepatocyte division. In addition to Fstl1, we identified Sfrp1 as a negative regulator that delays hepatocyte proliferation after PHx. Sfrp1, a Wnt pathway antagonist, has been reported as an inhibitor of liver cancer cell growth (Shih et al., 2007). Another study showed that Sfrp1 induces retinoblastoma senescence in vitro (Elzi et al., 2012).

Recently, single cell RNA-seq has been applied for the study of many liver processes and could also be applied to the study of liver regeneration (Halpern et al., 2017; MacParland et al., 2018). We chose the traditional method of bulk RNA sequencing for several reasons. First, genes for ligands and receptors are rarely expressed at high levels and the depth of the transcriptome is shallower with single cell data, especially for rare populations (Xiong et al., 2020). Second, we wish to provide a bulk RNA reference for single cell data generated by others. It will be interesting to see how well single cell RNA-seq captures non-abundant transcripts in rare cell populations.

In summary, we here provide a first-generation computational model of cell interactions within the mouse liver. Based on this model, we have characterized the contribution of all major cell types participating in the liver regeneration. Functional validation of a small subset of the interactions indicates that this network represents valuable real-life information on how the liver regenerates.

## Material and Methods

### Mice

Eight-week-old C57B/L6 male mice were purchased from The Jackson Laboratory. All animal experimentation was conducted in accordance with protocol IP00000445 of the institutional review committee at Oregon Health & Science University. To perform a 70% partial hepatectomy, the left lobe and median lobes were removed in the morning (Mitchell and Willenbring, 2008). For cell isolation after partial hepatectomy livers were perfused 24 hours after partial hepatectomy.

### Cell isolation and FACS sorting

To produce single liver cell suspensions for FACS, mouse livers were perfused with 0.5 mM EGTA (Fisher Scientific, O2783) followed by collagenase II (Worthington Biochemical). The isolation of hepatocytes and defined non-parenchymal cell (NPC) subpopulations from adult mouse liver was performed as previously described (Li et al., 2017; Li et al., 2019) with some modifications. Briefly, hepatocytes were pelleted at 50g for 5 min. The supernatant was then spun down at 1400 rpm for 5 min to pellet NPCs. Undigested tissue was further digested with collagenase IV (Worthington Biochemical) for 20 min at 37°C with stirring. Digested tissue was filtered with a 40µm cell strainer to collect NPCs. Leftover tissue was further digested with 0.25% Trypsin (Thermo Fisher) for another 20 min at 37°C with stirring. All digested mixtures were pooled to collect NPCs. For hepatocyte antibody labelling, cells were incubated at 4°C for 30 min with monoclonal anti OC2-2F8 hybridoma supernatant at a dilution of 1:20 and anti-CD45 at a concentration of 1:100. BEC, cBEC and NPD antibody labelling was performed as described (Li et al., 2017). Other antibodies used for FACS are listed in Table S5.

### Cell Culture

To test the effects of FSTL1 in vitro, hepatocytes were freshly isolated from 8 week old male C57B/L6 mice. Cryopreserved human hepatocytes were purchased from BioIVT. Both human and mouse hepatocytes were plated on collagen I coated tissue culture plate at a density of 1×10^5^/cm^2^ in William’E medium (Thermo Fisher) and 10% FBS (Thermo Fisher). Recombinant mouse Follistatin-like 1 protein (rFSTL1, R&D systems, MN, #1738-FN, 100ng/ml) was added into the culture media 24 h after plating cells and incubated for 24 hrs. After 24hrs, media was changed back to William’E media and 10% FBS without rFSTL1.

### RNA Sequencing

Cells were directly sorted into 0.5 ml TRIzol-LS (Thermo Fisher, 10296028) in 1.5 ml tubes (Eppendorf, CT) for RNA extraction. For rare sorted cell populations (<10,000 cells/mouse) multiple mice were perfused to collect and pool RNA. cDNA libraries were made with the Illumina TruSeq 2.0 (Illumina, CA) kit following the manufacturer’s instructions. The sequence reads were trimmed to 44 bases and aligned to the mouse genome NCBI37/mm9 using Bowtie (an ultrafast memory-efficient short-read aligner) version 0.12.7 (Langmead et al., 2009). We used custom scripts to count sequences in exons annotated for RefSeq mouse genes. DESeq2 (Love et al., 2014) was used to calculate the significance of differentially expressed genes based on these counts. We found some degree of Alb mRNA contamination in non-hepatocyte populations. To estimate hepatocyte contamination, we looked at a set of hepatocyte “marker” genes (“Serpina1a”, “Alb”, “Trf”, “Ttr”, “Hnf4a”, “Tat”, “Hpd”, “F9”, “Cyp2e1”, and “Cyp3a11”). For each of these genes, we calculated the average abundance in hepatocytes (based on FPKM values). We also calculated the abundance of these genes in each of the nonhepatocyte samples where these genes are not expected to be expressed.

Estimated contamination was based on the marker gene with the highest ratio in the test sample compared to the hepatocytes. To correct for this contamination, we calculated the average contribution of each gene in the hepatocytes and then applied the estimated contamination ratio from the sample being adjusted. For each gene, we subtracted this value. Finally, we updated the FPKM values of the adjusted sample to account for the loss in overall expression level. This process was applied separately for pre and post partial hepatectomy samples. The RNA-seq FASTQ data were submitted to the NCBI GSE226004.

### CCInx analyses

The algorithm was adapted from previous analyses (Qiao et al., 2014). To find active receptor-ligand interactions, we used the distribution of expression levels in our data to select an “On” (active) level for each gene. Specifically, we applied a single factor ANOVA test to each gene with the factor being cell type (pre and post partial hepatectomy were handled separately). For genes with significant (p-value <= 0.01) changes in expression, we divided into two groups by k-means clustering and defined these as “On” (higher expression) and “Off” (lower expression) states. For each cell type, we checked to see if a gene is called as “On” for all replicates and if the average expression of all replicates is at least 2-fold greater than the average expression of all cases of this gene in the “Off” state (averaging is done on log scaled data). If so, this gene is called as “On” for that cell type. A ligand/receptor interaction is only considered valid when both associated genes are “On” for the respective cell types. The magnitude of each interaction is based on the average log2 fold change of the “On” states for the respective cell types. However, this log2 fold change value is set to zero for genes that are not called as “On”.

### Plasmids

The *Fstl1* cDNA clone was purchased from GeneCopoeia Mm02579, NCBI entry NM_008047. To make AAV-Fstl1, a DNA fragment containing the human thyroxine binding globulin (TBG) promoter (Yan et al., 2012) and Fstl1 transgene were cloned into a self-complementary rAAV vector between two inverted terminal repeats (ITRs). The primers for In-fusion cloning (Takara Bio) Fstl1 are (Forward 5’-CACAGACGCGTACCGGTGCCACCATGTGGAAACGATGGCTGGCGCTC-3’, Reverse 5’-GTGAGGCCTAGCGGCCGCTTAGATCTCTTTGGTGTTCACCTT-3’).

Mouse cDNA was synthesized by reverse transcription from kidney RNA by M-MLV (ThermoFisher 28025013) with random primers (Invitrogen 48190011). A full-length *Sfrp1* cDNA fragment was obtained by PCR amplifying mouse kidney cDNA using PrimeStar HS DNA polymerase (Takara Bio) following manufacturer’s protocol. The primers for Sfrp1 In-fusion cloning (Takara Bio) are (Forward 5’-CTAGTGATT TCGCCGCCACCATGGGCGTCGGGCGCAGCGCGCG-3’, Reverse 5’-TCAGGTCAGCTATCTTACTTGTACAAGCTTCACTTAAAAACAGACTGGAAGGTG-3’). To make AAV-Sfrp1, the DNA fragment containing a liver specific promoter1 (LSP1) (A gift from Hiroyuki Nakai lab) and *Sfrp1* transgene was cloned into a self-complementary AAV vector between two ITRs.

Sequences for Fstl1, Sfrp1 cDNAs and TBG promoter are listed in Table S6.

### rAAV production and *in vivo* administration

Recombinant Fstl1 was packaged into the AAV-DJ capsid (Grimm et al., 2008) and Sfrp1 was produced with serotype AAV8 (Gao et al., 2002). The rAAV vector preps were made using the standard triple plasmid co-transfection method and purified with Iodixanol Gradient Ultracentrifugation (Zhang et al., 2021). rAAV titers were determined by dot-blot hybridization (Zhang et al., 2021). For AAVDJ-Fstl1 administration, 8-week-old male mice obtained from Jackson Laboratory received 2×10^11^ vg of AAVDJ-Fstl1 or AAVDJ-tdTomato (control) vector diluted in saline solution (100 µl total) via retro-orbital injection. For AAV8-Sfrp1 administration, 3 and 8-week-old male mice obtained from Jackson Laboratory received 2×10^11^ vg of AAV8-SFRP1 or AAV8-GFP (control) vector diluted in saline solution (100 µl total) via retro-orbital injection. One week after rAAV injection, mice were subjected to 70% partial hepatectomy and sacrificed as indicated.

### BrdU labeling

BrdU (Thermo Fisher Scientific, H27260) was given in the drinking water (0.5 mg/mL) as indicated. For BrdU histochemistry, the sections were treated with 2N HCl for 1 hr and then stained with anti-BrdU (Abcam, ab6326) antibody (Willenbring et al., 2008).

### Statistical Analyses

All data are presented as mean ± SD. GraphPad Prism software was used for statistical analyses. p < 0.05 and p < 0.01 were considered to be statistically significant and highly significant, respectively.

## Accession Numbers

The accession number for the RNA-seq data reported in this paper is GSE226004.

## Author contributions

M.G. supervised, designed the experiments and wrote the manuscript. B.L. and D.R-T. designed and conducted the experiments, and wrote the manuscript. C.P. designed and generated the RNA-seq analysis data and adapted CCInx. B.I., P.Z. and G.B. generated the cell network platform. S.C. assisted with vector construction. P.C. assisted with the flow cytometry-related experiments.

## Acknowledgment

AAV plasmids with TBG and LSP1 promoters were generous gifts from the Hiroyuki Nakai lab. This work was supported by NIH grants R01-DK051592 and R01DK083355 to M.G. DR-T was supported by the Alfonso Martín Escudero Foundation Fellowship. The authors thank OHSU core services Massively Parallel Sequencing Shared Resource and OHSU Flow Cytometry Shared Resources. We thank Milton Finegold and Angela Major for histology assistance. We also thank Leslie Wakefield, Geoff Clarke, Shinichiro Ogawa, and Jeffrey Posey for their excellent technical assistances. M.G. is a founder and shareholder of Yecuris and Ambys Medicine.

## Abbreviations

AAV: adeno-associated virus BEC: biliary epithelial cell
BrdU: 5-bromo-2’-deoxyuridine cBEC: clonal biliary epithelial cell
CCInx: Cell-cell interactions EC: endothelial cell
HC: hepatocyte
HSC: hepatic stellate cell
LSEC: liver sinusoidal endothelial cell NPC: non-parenchymal cell
NPD: non-progenitor duct cell PHx: partial hepatectomy

## References

1. Alaverdi, N. (2004). Monoclonal antibodies to mouse cell-surface antigens. Current protocols in immunology 62, A. 4B. 1-A. 4B. 25.

2. Azuma, H., Paulk, N., Ranade, A., Dorrell, C., Al-Dhalimy, M., Ellis, E., Strom, S., Kay, M.A., Finegold, M., and Grompe, M. (2007). Robust expansion of human hepatocytes in Fah−/−/Rag2−/−/Il2rg−/− mice. Nature biotechnology 25, 903–910.

3. Ding, B.-S., Nolan, D.J., Butler, J.M., James, D., Babazadeh, A.O., Rosenwaks, Z., Mittal, V., Kobayashi, H., Shido, K., and Lyden, D. (2010). Inductive angiocrine signals from sinusoidal endothelium are required for liver regeneration. Nature 468, 310–315.

4. Dorrell, C., Erker, L., Schug, J., Kopp, J.L., Canaday, P.S., Fox, A.J., Smirnova, O., Duncan, A.W., Finegold, M.J., and Sander, M. (2011). Prospective isolation of a bipotential clonogenic liver progenitor cell in adult mice. Genes & development 25, 1193–1203.

5. Elzi, D.J., Song, M., Hakala, K., Weintraub, S.T., and Shiio, Y. (2012). Wnt antagonist SFRP1 functions as a secreted mediator of senescence. Molecular and cellular biology 32, 4388–4399.

6. Gao, G.P., Alvira, M.R., Wang, L., Calcedo, R., Johnston, J., and Wilson, J.M. (2002). Novel adenoassociated viruses from rhesus monkeys as vectors for human gene therapy. Proc Natl Acad Sci U S A 99, 11854–11859.

7. Geng, Y., Dong, Y., Yu, M., Zhang, L., Yan, X., Sun, J., Qiao, L., Geng, H., Nakajima, M., and Furuichi, T. (2011). Follistatin-like 1 (Fstl1) is a bone morphogenetic protein (BMP) 4 signaling antagonist in controlling mouse lung development. Proceedings of the National Academy of Sciences 108, 7058–7063.

8. Grimm, D., Lee, J.S., Wang, L., Desai, T., Akache, B., Storm, T.A., and Kay, M.A. (2008). In vitro and in vivo gene therapy vector evolution via multispecies interbreeding and retargeting of adeno-associated viruses. J Virol 82, 5887–5911.

9. Halpern, K.B., Shenhav, R., Matcovitch-Natan, O., Toth, B., Lemze, D., Golan, M., Massasa, E.E., Baydatch, S., Landen, S., and Moor, A.E. (2017). Single-cell spatial reconstruction reveals global division of labour in the mammalian liver. Nature 542, 352–356.

10. Hoppo, T., Fujii, H., Hirose, T., Yasuchika, K., Azuma, H., Baba, S., Naito, M., Machimoto, T., and Ikai, I. (2004). Thy1-positive mesenchymal cells promote the maturation of CD49f-positive hepatic progenitor cells in the mouse fetal liver. Hepatology 39, 1362–1370.

11. Huch, M., Dorrell, C., Boj, S.F., Van Es, J.H., Li, V.S., Van De Wetering, M., Sato, T., Hamer, K., Sasaki, N., and Finegold, M.J. (2013). In vitro expansion of single Lgr5+ liver stem cells induced by Wnt-driven regeneration. Nature 494, 247–250.

12. Huh, C.G., Factor, V.M., Sanchez, A., Uchida, K., Conner, E.A., and Thorgeirsson, S.S. (2004). Hepatocyte growth factor/c-met signaling pathway is required for efficient liver regeneration and repair. Proc Natl Acad Sci U S A 101, 4477–4482.

13. Innes, B.T., and Bader, G.D. (2021). Transcriptional signatures of cell-cell interactions are dependent on cellular context. bioRxiv.

14. Ishii, T., Sato, M., Sudo, K., Suzuki, M., Nakai, H., Hishida, T., Niwa, T., Umezu, K., and Yuasa, S. (1995). Hepatocyte growth factor stimulates liver regeneration and elevates blood protein level in normal and partially hepatectomized rats. J Biochem 117, 1105–1112.

15. Kaibori, M., Inoue, T., Oda, M., Naka, D., Kawaguchi, T., Kitamura, N., Miyazawa, K., Kwon, A.H., Kamiyama, Y., and Okumura, T. (2002). Exogenously administered HGF activator augments liver regeneration through the production of biologically active HGF. Biochem Biophys Res Commun 290, 475–481.

16. Kirouac, D.C., Ito, C., Csaszar, E., Roch, A., Yu, M., Sykes, E.A., Bader, G.D., and Zandstra, P.W. (2010). Dynamic interaction networks in a hierarchically organized tissue. Molecular systems biology 6, 417.

17. Kirouac, D.C., Madlambayan, G.J., Yu, M., Sykes, E.A., Ito, C., and Zandstra, P.W. (2009). Cell–cell interaction networks regulate blood stem and progenitor cell fate. Molecular systems biology 5, 293.

18. Kordes, C., Sawitza, I., Gotze, S., Herebian, D., and Haussinger, D. (2014). Hepatic stellate cells contribute to progenitor cells and liver regeneration. J Clin Invest 124, 5503–5515.

19. Kumar, V., Ali, S.R., Konrad, S., Zwirner, J., Verbeek, J.S., Schmidt, R.E., and Gessner, J.E. (2006). Cellderived anaphylatoxins as key mediators of antibody-dependent type II autoimmunity in mice. The Journal of clinical investigation 116, 512–520.

20. Langmead, B., Trapnell, C., Pop, M., and Salzberg, S.L. (2009). Ultrafast and memory-efficient alignment of short DNA sequences to the human genome. Genome biology 10, 1–10.

21. Li, B., Dorrell, C., Canaday, P.S., Pelz, C., Haft, A., Finegold, M., and Grompe, M. (2017). Adult mouse liver contains two distinct populations of cholangiocytes. Stem cell reports 9, 478–489.

22. Li, B., Dorrell, C., Canady, P.S., and Wakefield, L. (2019). Identification and isolation of clonogenic cholangiocyte in mouse. In Hepatic Stem Cells (Springer), pp. 19–27.

23. Li, Y., Fan, W., Link, F., Wang, S., and Dooley, S. (2022). Transforming growth factor beta latency: A mechanism of cytokine storage and signalling regulation in liver homeostasis and disease. JHEP Rep 4, 100397.

24. Love, M.I., Huber, W., and Anders, S. (2014). Moderated estimation of fold change and dispersion for RNA-seq data with DESeq2. Genome biology 15, 1–21.

25. MacParland, S.A., Liu, J.C., Ma, X.-Z., Innes, B.T., Bartczak, A.M., Gage, B.K., Manuel, J., Khuu, N., Echeverri, J., and Linares, I. (2018). Single cell RNA sequencing of human liver reveals distinct intrahepatic macrophage populations. Nature communications 9, 1–21.

26. Mead, J.E., and Fausto, N. (1989). Transforming growth factor alpha may be a physiological regulator of liver regeneration by means of an autocrine mechanism. Proc Natl Acad Sci U S A 86, 1558–1562.

27. Mederacke, I., Dapito, D.H., Affò, S., Uchinami, H., and Schwabe, R.F. (2015). High-yield and high-purity isolation of hepatic stellate cells from normal and fibrotic mouse livers. Nature protocols 10, 305–315.

28. Michalopoulos, G.K. (2007). Liver regeneration. Journal of cellular physiology 213, 286-300.

29. Michalopoulos, G.K., and DeFrances, M.C. (1997). Liver regeneration. Science 276, 60–66.

30. Mitchell, C., and Willenbring, H. (2008). A reproducible and well-tolerated method for 2/3 partial hepatectomy in mice. Nat Protoc 3, 1167-1170.

31. Nejak-Bowen, K., Orr, A., Bowen, W.C., Jr., and Michalopoulos, G.K. (2013). Conditional genetic elimination of hepatocyte growth factor in mice compromises liver regeneration after partial hepatectomy. PLoS One 8, e59836.

32. Patijn, G.A., Lieber, A., Schowalter, D.B., Schwall, R., and Kay, M.A. (1998). Hepatocyte growth factor induces hepatocyte proliferation in vivo and allows for efficient retroviral-mediated gene transfer in mice. Hepatology 28, 707–716.

33. Qiao, W., Wang, W., Laurenti, E., Turinsky, A.L., Wodak, S.J., Bader, G.D., Dick, J.E., and Zandstra, P.W. (2014). Intercellular network structure and regulatory motifs in the human hematopoietic system. Mol Syst Biol 10, 741.

34. Russell, J.O., and Monga, S.P. (2018). Wnt/β-catenin signaling in liver development, homeostasis, and pathobiology. Annual Review of Pathology: Mechanisms of Disease 13, 351–378.

35. Shih, Y.L., Hsieh, C.B., Lai, H.C., Yan, M.D., Hsieh, T.Y., Chao, Y.C., and Lin, Y.W. (2007). SFRP1 suppressed hepatoma cells growth through Wnt canonical signaling pathway. International journal of cancer 121, 1028–1035.

36. Shinozuka, H., Kubo, Y., Katyal, S.L., Coni, P., Ledda-Columbano, G.M., Columbano, A., and Nakamura, T. (1994). Roles of growth factors and of tumor necrosis factor-alpha on liver cell proliferation induced in rats by lead nitrate. Laboratory investigation; a journal of technical methods and pathology 71, 35–41.

37. Vrochides, D., Papanikolaou, V., Pertoft, H., Antoniades, A.A., and Heldin, P. (1996). Biosynthesis and degradation of hyaluronan by nonparenchymal liver cells during liver regeneration. Hepatology 23, 1650–1655.

38. Wei, K., Serpooshan, V., Hurtado, C., Diez-Cunado, M., Zhao, M., Maruyama, S., Zhu, W., Fajardo, G., Noseda, M., and Nakamura, K. (2015). Epicardial FSTL1 reconstitution regenerates the adult mammalian heart. Nature 525, 479–485.

39. Willenbring, H., Sharma, A.D., Vogel, A., Lee, A.Y., Rothfuss, A., Wang, Z., Finegold, M., and Grompe, M. (2008). Loss of p21 permits carcinogenesis from chronically damaged liver and kidney epithelial cells despite unchecked apoptosis. Cancer Cell 14, 59–67.

40. Ximerakis, M., Lipnick, S.L., Innes, B.T., Simmons, S.K., Adiconis, X., Dionne, D., Mayweather, B.A., Nguyen, L., Niziolek, Z., and Ozek, C. (2019). Single-cell transcriptomic profiling of the aging mouse brain. Nature neuroscience 22, 1696–1708.

41. Xiong, X., Kuang, H., Liu, T., and Lin, J.D. (2020). A Single-Cell Perspective of the Mammalian Liver in Health and Disease. Hepatology 71, 1467–1473.

42. Yan, Z., Yan, H., and Ou, H. (2012). Human thyroxine binding globulin (TBG) promoter directs efficient and sustaining transgene expression in liver-specific pattern. Gene 506, 289–294.

43. Zaret, K.S., and Grompe, M. (2008). Generation and regeneration of cells of the liver and pancreas. Science 322, 1490–1494.

44. Zhang, Q.-S., Tiyaboonchai, A., Nygaard, S., Baradar, K., Major, A., Balaji, N., and Grompe, M. (2021). Induced liver regeneration enhances CRISPR/Cas9-mediated gene repair in tyrosinemia type 1. Human Gene Therapy 32, 294–301.

