## Supplemental Figures for "Cell networks in the mouse liver during partial hepatectomy"

**Figure S1 FACS strategy for isolating liver cells**

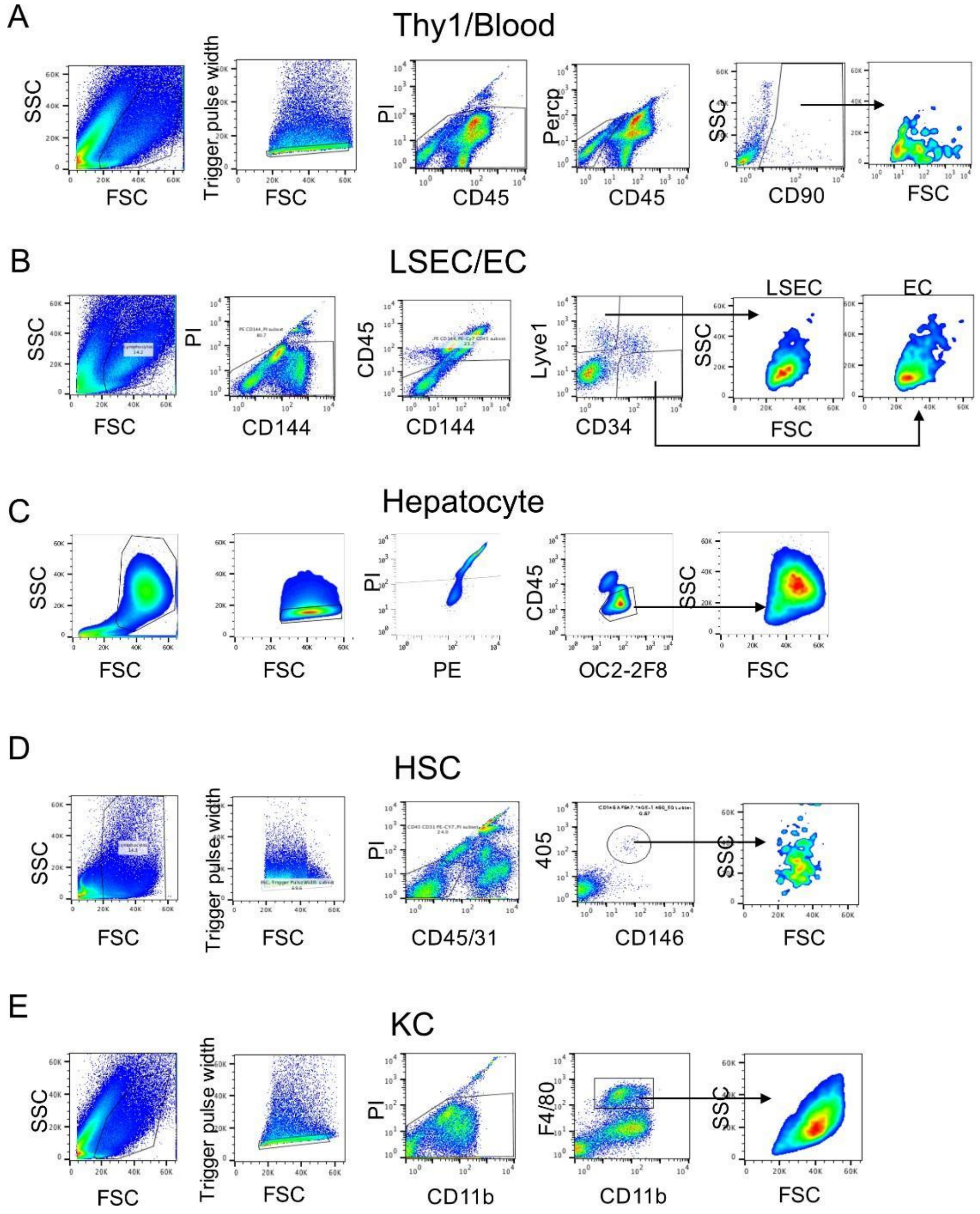


Single cell suspension was labelled with antibodies to isolate liver cell populations

Cells were sequentially gated based on cell size (forward scatter [FSC] versus side scatter [SSC]) and singlets (FSC versus trigger pulse width). Dead cells and debris were gated out with propidium iodide (PI) positivity. (A) Blood cells were isolated with CD45 positivity and Thy1 cells were isolated with $CD45^{-}CD90^{+}$. (B) LSEC and EC were isolated with $CD45^{-}$

CD144$^{+}$Lyve1$^{+}$CD34$^{-}$ and CD45$^{-}$CD144$^{-}$Lyve1$^{+}$CD34$^{+}$. (C) Hepatocytes were first gated with bigger FSC vs SSC, singlets followed by CD45$^{-}$OC2-2F8$^{+}$. (D) Hepatic stellate cells (HSC) were isolated by CD45$^{-}$CD31$^{-}$Violet$^{+}$CD146$^{+}$. (E) Kupffer cells (KC) were isolated with CD11b$^{+}$F4/80$^{+}$.

**Figure S2 Ligand-receptor analysis by CCInx**

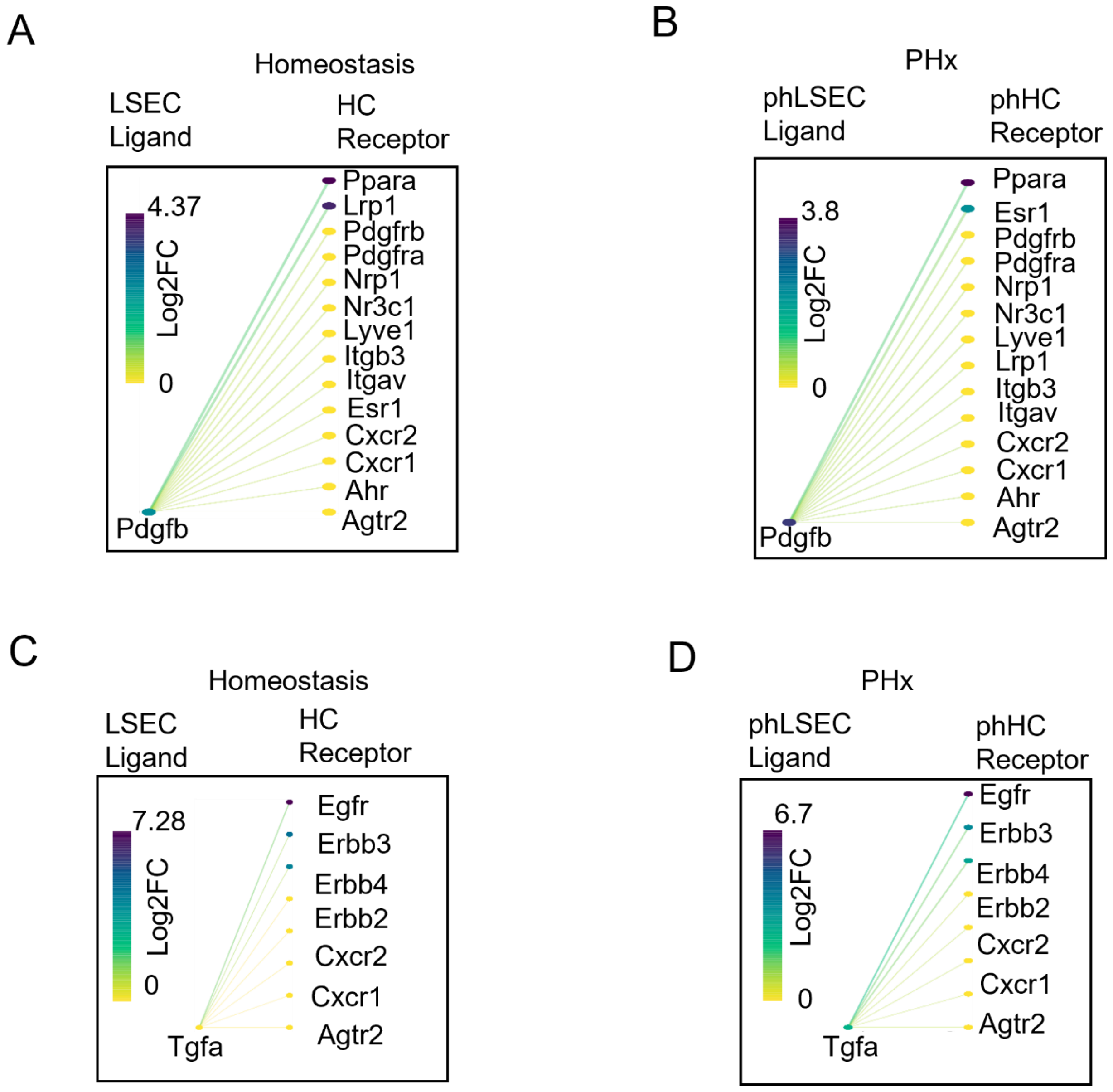


(A-B) Pdgfβ ligand in LSEC and its receptor in hepatocyte in normal (A) and liver regeneration (B), respectively.

(C-D) Tgfα ligand in LSEC and its receptor in hepatocyte in normal (C) and liver regeneration (D), respectively. Log2FC: log2fold change.

**Figure S3 Effects of rFstl1 on mouse and human hepatocytes**

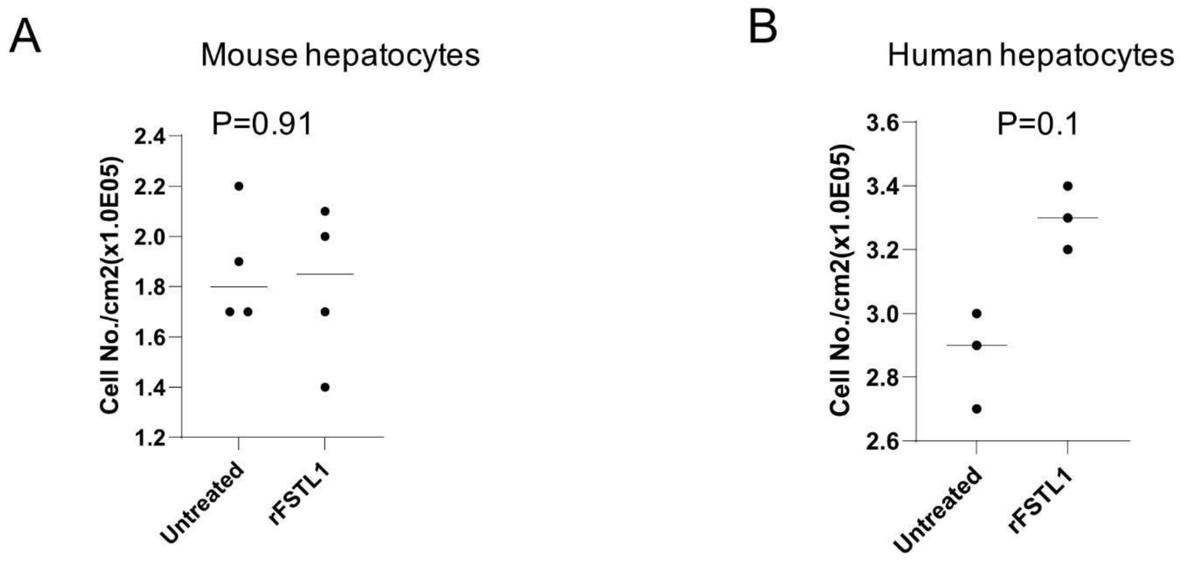


Recombinant FSTL1(rFSTL1) was added into the cell culture supernatant.

(A) Cell densities for mouse hepatocytes culture treated with rFSTL1 and untreated ($n = 4$ for each treatment). Statistical analyses: Student's t test, $p = 0.91$.

(B) Cell densities for human hepatocytes cultured with rFSTL1 and untreated ($n = 3$). Statistical analyses: Student's t test, $p = 0.1$.

### Table S1 OnOn interactions

This table summarizes the numbers of receptor/ligand interactions. Each sheet reports numbers for all interactions and numbers for interactions involving selected genes. The first sheet reports counts for interactions between all cell types. The additional sheets report interactions involving each cell type and involving each cell type specifically as a ligand or as a receptor. The columns for each sheet are: Gene - either all genes or one of a list of selected genes of interest, ValidInteractionPairs - number of possible interactions, OnOnInGroundState - number of active interactions in the pre partial hepatectomy state (pre PHx), OnOnInPHxState - number of active interactions in the post partial hepatectomy state (post PHx), OnOnInBothStates - number of active interactions shared in both states, OnOnInGroundStateOnly - number of active interactions specific to the pre PHx state, and OnOnInPHxStateOnly - number of active interactions specific to the post PHx state. To determine OnOn interactions we first estimated an on state expression level threshold for each gene based on the distribution of expression levels in our data (this was done independently in the pre and post PHx states, see methods). We then required that all replicates of respective cell types exceed this threshold for a receptor/ligand pair in the database. Only these cases are considered valid OnOn interactions.

### Table S2 All of the interactions

This table shows all active interactions between all cell types. Active interactions are reported for pre and post partial hepatectomy states separately. The columns are: Name - the format is gene1_cell type1~gene2_cell type2 where the cell type 1 is interacting with cell type 2 via gene 1 and gene 2 respectively (the receptor/ligand order is not defined), GeneStateLog2FCnodeA - the log2 Fold Change (FC) for node A (gene 1, cell type 1), GeneStateLog2FCnodeB - the log2 FC for node B (gene 2, cell type 2), Rank - the rank order of these interactions based on the maximum of the two log2 FC values. Note that the log2 FC is calculated based on expression in the given cell type compared to the off state expression level estimated based on all cell types (see methods).

### Table S3 Top10 weighted interactions in normal liver by CCInx

This table shows the top active interactions in normal liver for all cell types. The top 10 CCInx weighted interactions from each cell type are combined (duplicates are removed). The columns are: Ligand_Cell_Type - Cell type expressing ligand, Receptor_Cell_Type - Cell type expressing receptor, Interaction_Type - Paracrine or Autocrine, Name - Formatted name of interaction, Interaction_Weight - Relative strength of interaction from CCInx tool, Ligand_log2FC - Log 2 Fold Change (FC) of ligand relative to calculated off state in respective cell type, Ligand_Symbol - Gene symbol of ligand, Ligand_Weight - Relative degree of ligand expression from CCInx tool, Receptor_log2FC - Log 2 FC of receptor relative to calculated off state in respective cell type, Receptor_Symbol - Gene symbol of receptor, Receptor_Weight - Relative degree of receptor expression from CCInx tool.

### Table S4 Top10 weighted interactions in PHx liver by CCInx

This table shows the top active interactions in PHx liver for all cell types. The top 10 CCInx weighted interactions from each cell type are combined (duplicates are removed). The columns are: Ligand_Cell_Type - Cell type expressing ligand, Receptor_Cell_Type - Cell type expressing receptor, Interaction_Type - Paracrine or Autocrine, Name - Formatted name of interaction, Interaction_Weight - Relative strength of interaction from CCInx tool, Ligand_log2FC - Log 2 Fold Change (FC) of ligand relative to calculated off state in respective cell type, Ligand_Symbol - Gene symbol of ligand, Ligand_Weight - Relative degree of ligand expression from CCInx tool, Receptor_log2FC - Log 2 FC of receptor relative to calculated off state in respective cell type, Receptor_Symbol - Gene symbol of receptor, Receptor_Weight - Relative degree of receptor expression from CCInx tool.

### Table S5 List of Antibodies

Primary Antibodies

| Name | IgG type | Use | Dilution | Company | Lot |
|---|---|---|---|---|---|
| CD11b | Rat mAB | FACS | 1/100 | BD biosciences | 552850 |
| CD144 | Rat mAB | FACS | 1/100 | eBioscience | 12-1441-82 |
| CD146 | Rat mAB | FACS | 1/100 | BD biosciences | 562230 |
| CD26 | Rat mAB | FACS | 1/100 | BD biosciences | 4278805 |
| CD309 | Biotin | FACS | 1/100 | eBioscience | 13-5821-82 |
| CD31 | Rat mAB | FACS | 1/100 | BD biosciences | 561410 |
| CD34 | Rat mAB | FACS | 1/20 | eBioscience | 50-0341-80 |
| CD45 | Rat mAB | FACS | 1/100 | BD biosciences | 552848 |

| | | | | | |
|---|---|---|---|---|---|
| CD90(Thy1) | Rat mAB | FACS | 1/100 | BD biosciences | 553007 |
| F4/80 | Rat mAB | FACS | 1/100 | eBioscience | 17-4801-80 |
| Lyve1 | Rat mAB | FACS | 1/100 | eBioscience | 53-0443-80 |
| MIC1-1C3 | Rat mAB | FACS | 1/20 | Grompe lab | |
| OC2-2F8 | Rat mAB | FACS | 1/20 | Grompe lab | |
| ST14 | Rabbit pAB | FACS | 1/100 | Abcam | ab28266 |
| BrdU | Rat mAB | IHC | 1/100 | Abcam | Ab6326 |

Secondary Antibodies

| Name | IgG type | Use | Dilution | Company | Lot |
|---|---|---|---|---|---|
| Anti rabbit APC | Donkey | FACS | 1/200 | JacksonImmuno | 711-056-152 |
| Anti Rat PE | Goat | FACS | 1/200 | JacksonImmuno | 112-116-143 |
| APC/Cy7 | Streptavidin | FACS | 1/250 | Biolegend | 405208 |
| Goat Anti-Rat | HRP | IHC | 1/500 | JacksonImmuno | 112-035-003 |

**Table S6 Plasmid Sequence**

| | |
|---|---|
| Fstl1 | ATGTGGAAACGATGGCTGGCGCTCTCGCTGGTGACCATCGCCCTGGTCCACGG CGAGGAGGAACCTAGAAGCAAATCCAAGATCTGCGCCAATGTGTTTTGTGGAGC TGGCAGGGAATGTGCCGTCACAGAGAAGGGGGAGCCCACGTGCCTCTGCATTG AGCAATGCAAACCTCACAAGAGGCCTGTGTGTGGCAGTAATGGCAAGACCTACC TCAACCACTGTGAACTTCATAGAGATGCCTGCCTCACTGGATCCAAGATCCAGGT TGATTATGATGGGCACTGCAAAGAAAAGAAGTCTGCGAGTCCATCTGCCAGCCC AGTTGTCTGCTATCAAGCTAACCGCGATGAGCTCCGACGGCGCCTCATCCAGTG GCTGGAAGCTGAGATCATTCCAGATGGCTGGTTCTCTAAAGGCAGTAACTACAGT GAGATCCTAGACAAGTACTTTAAGAGCTTTGATAATGGCGACTCTCACCTGGACT CCAGTGAATTCCTGAAATTCGTGGAGCAGAATGAAACAGCCATCAACATCACCAC TTATGCAGATCAGGAGAACAACAAACTGCTCAGAAGCCTCTGTGTTGACGCCCTC ATTGAACTGTCTGATGAGAACGCTGACTGGAAACTCAGCTTCCAAGAGTTCCTCA AGTGCCTCAACCCATCCTTCAACCCTCCTGAGAAGAAGTGTGCCCTGGAGGACG AAACCTATGCAGATGGAGCTGAGACTGAGGTGGACTGCAATCGCTGTGTCTGTT CCTGTGGCCACTGGGTCTGCACAGCAATGACCTGTGATGGAAAGAATCAGAAGG GGGTCCAGACCCACACAGAGGAGGAGAAGACAGGATATGTCCAGGAACTCCAG AAGCACCAGGGCACAGCAGAAAAGACCAAGAAGGTGAACACCAAAGAGATCTAA |
| TBG | TGCATGTATAATTTCTACAGAACCTATTAGAAAGGATCACCCAGCCTCTGCTTTTG TACAACTTTCCCTTAAAAAACTGCCAATTCCACTGCTGTTTGGCCCAATAGTGAGA ACTTTTTCCTGCTGCCTCTTGGTGCTTTTGCCTATGGCCCCTATTCTGCCTGCTG AAGACACTCTTGCCAGCATGGACTTAAACCCCTCCAGCTCTGACAATCCTCTTTC TCTTTTGTTTTACATGAAGGGTCTGGCAGCCAAAGCAATCACTCAAAGTTCAAAC CTTATCATTTTTTGCTTTGTTCCTCTTGGCCTTGGTTTTGTACATCAGCTTTGAAAA TACCATCCCAGGGTTAATGCTGGGGTTAATTTATAACTAAGAGTGCTCTAGTTTTG CAATACAGGACATGCTATAA |
| Sfrp1 | ATGGGCGTCGGGCGCAGCGCGCGGGGTCGCGGCGGGGCCGCCTCGGGAGTG CTGCTGGCGTTGGCCGCCGCTCTGCTGGCCGCGGGTTCGGCCAGCGAGTACGA CTACGTGAGCTTCCAGTCCGACATCGGCTCGTATCAGAGCGGGCGCTTCTACAC CAAGCCCCCGCAGTGCGTGGACATCCCGGTGGACCTGAGGCTGTGCCACAACG TGGGCTACAAGAAGATGGTGCTGCCCAACCTGCTGGAGCACGAGACCATGGCA GAGGTGAAGCAGCAGGCCAGCAGCTGGGTGCCGCTGCTCAACAAGAACTGCCA CATGGGCACCCAGGTCTTCCTCTGTTCGCTCTTCGCGCCCGTCTGTCTGGACCG GCCCATCTACCCGTGTCGCTGGCTCTGCGAGGCCGTGCGCGACTCGTGCGAGC CGGTCATGCAGTTCTTCGGCTTCTACTGGCCCGAGATGCTCAAATGTGACAAGTT CCCCGAGGGCGACGTCTGCATCGCCATGACCCCGCCCAATACCACGGAAGCCT CTAAGCCCCAAGGTACAACCGTGTGTCCTCCATGCGACAACGAGTTGAAGTCAG AGGCCATCATTGAACATCTCTGTGCAAGCGAGTTTGCACTGAGGATGAAAATCAA AGAAGTGAAGAAGGAAAACGGTGACAAGAAGATTGTCCCCAAGAAGAAGAAACC CTTGAAGCTGGGGCCCATCAAGAAGAAGGAGCTGAAGCGGCTTGTGCTGTTCCT GAAGAACGGTGCCGACTGTCCCTGCCACCAGCTGGACAACCTCAGCCACAACTT TCTCATCATGGGCCGCAAGGTGAAGAGCCAGTACCTGCTGACAGCCATTCACAA GTGGGACAAGAAAAACAAGGAGTTCAAAAACTTCATGAAGAGAATGAAAAACCAC GAGTGTCCCACCTTCCAGTCTGTTTTTAAGTGAAGCTTGTACAAGTAA |
